# Brain-wide connectivity and novelty response of the dorsal endopiriform nucleus in mice

**DOI:** 10.1101/2024.09.30.615899

**Authors:** Steffy B. Manjila, Seoyoung Son, Hannah Kline, Rebecca Betty, Yuan-ting Wu, Pi Hyun-Jae, Deniz Parmaksiz, Donghui Shin, Josephine K. Liwang, Fae N. Kronman, Anirban Paul, Yongsoo Kim

**Affiliations:** Department of Neural and Behavioral Sciences, The Pennsylvania State University, Hershey, PA, 17033, USA; Department of Pharmacology, School of Medicine, Wayne State University, MI 48201; Department of Neurosurgery, Department of Computational Biomedicine, Cedars-Sinai Medical Center, CA 90048

**Keywords:** dorsal endopiriform nucleus, whole brain mapping, brain connectivity, spatial transcriptome, oxytocin receptor, exploratory behavior

## Abstract

The dorsal endopiriform nucleus (EPd) is an enigmatic cortical subplate structure located inside the piriform cortex that shares a similar developmental origin with the claustrum. Although the EPd has been previously implicated in epilepsy and olfactory processing, its anatomical organization, connectivity patterns, and function remain largely unclear due to a lack of specific molecular markers. Our previous mapping study serendipitously identified that the Oxt receptor (Oxtr) is densely expressed in the EPd. Subsequent immunohistochemical and spatial transcriptomic analyses confirmed that Oxtr expression is enriched in the EPd, revealing distinct molecular organization compared to the neighboring claustrum. Whole brain input-output mapping of EPd Oxtr-positive neurons unveils extensive bidirectional connections to the ventral half of the brain, orchestrating functional circuits regulating olfaction, internal state, and emotion. Furthermore, our in-vivo miniscope recordings show that EPd Oxtr neurons exhibit high baseline activity during exploratory behavior, with a sharp decrease in activity in response to novel stimuli. This suggests that the EPd regulates interoceptive states and likely plays a role in adapting to novel exteroceptive cues.

## Introduction

The dorsal endopiriform nucleus (EPd) is a thin, elongated structure along the anterior-posterior axis in the mouse brain that is situated medially (endo-) to the piriform cortex(*1*, *2*). Moreover, the EPd is located underneath the claustrum (CLA) in the dorsoventral axis and shares a similar developmental origin with the CLA(*3*). The EPd is a conserved structure in rodent brains and higher mammals and is a part of the claustrum complex(*4*). While the CLA forms extensive connections with iso (neo)cortical areas(*5–9*), previous studies suggest that the EPd exhibits widespread connectivity with the olfactory areas and limbic cortices(*2*, *10–12*). Although the functions of the EPd are largely unknown, a few studies have demonstrated its potential involvement in multisensory (olfactory and gustatory)(*12*), limbic information processing(*2*), and conscious perception(*13*). Moreover, dysfunction of the claustrum complex including the EPd has been implicated in many neurological conditions such as epilepsy, autism, and neurodegenerative disorders(*14–17*). Despite the potential significance of the EPd, very little information is known about this region, including basic neuroanatomy, molecular and cellular characteristics, and its function in regulating behavior.

One of the major challenges in examining the EPd is the lack of clear molecular markers for targeted studies in the area. While we investigated Oxt receptor (Oxtr) expression throughout the whole mouse brain(*18*), we serendipitously found that Oxtr is densely expressed in the EPd. In this study, we utilized in situ hybridization and spatial transcriptomics and confirmed that Oxtr is indeed richly expressed in the EPd. This offers new opportunities to perform initial groundwork to characterize the cellular architecture of the EPd, its connectivity and potential functions.

Detailed anatomical wiring diagrams have been crucial to gaining accurate anatomical connectivity of target areas, which helps to infer relevant function. For instance, our Oxt wiring diagram helps to identify nine functional circuits where Oxt signal can modulate different functions(*19*). Here, we perform detailed anatomical studies to investigate the input-output connectivity of the EPd using Oxtr-Cre mice. Our findings unveil extensive bidirectional connections with ventral brain regions, particularly limbic and olfactory areas, in stark contrast with neighboring CLA connectivity patterns largely targeting the dorsal brain regions including the isocortex (*5*, *7*, *20*, *21*). Moreover, we found unidirectional mono-synaptic input from specific thalamic, midbrain, and hindbrain areas that are linked with the limbic system and alertness. Lastly, our results show that EPd neurons exhibit high baseline neural activity during exploratory behavior, and significantly decreases their activities when exposed to novel stimuli, including novel social exposure. Collectively, our findings imply that the EPd is well-positioned to regulate behavior driven by interoception vs. exteroception

## Results

### Molecular signature of the dorsal endopiriform nucleus (EPd) in the mouse brain

Cell types with distinct gene expression patterns have been crucial in defining specific anatomical structures(*22–26*). To identify shared and distinct gene expression signatures in the EPd against neighboring areas, we performed Multiplexed Error-Robust Fluorescence in situ Hybridization (MERFISH) based spatial transcriptomics using known cell type and area-specific markers such as Nr4a2 (also called Nurr1)(*27–29*), which help to identify the EPd, the CLA, and the neighboring cortical areas(*30*)(Figure 1A-B). We performed unbiased gene clustering analysis with known cell type markers and found 12 different cell type clusters(*31*) (Figure 1C). When anatomical areas are overlaid in the UMAP, we found that the CLA and the EPd exist in similar clusters, distinct from neighboring piriform cortex (PIR) and the agranular insular cortex (AI) (Figure 1D). Then, we examined genes differentially enriched in the CLA-EPd area. We found that Ccn2, Nxph3, Npsr1, Nxph4 and Ccdc80 are more significantly expressed in the EPd (Figure 1E) while Satb2, Bmp3, Cadps2, Gnb4, and Gng2 are enriched in the CLA (Supplementary Figure 1A). The AI and PIR showed different sets of genes enriched compared to the CLA-EPd (Supplementary Figure 1B-C), further supporting that the EPd has a distinct gene expression pattern compared to neighboring areas.

**Figure 1.**
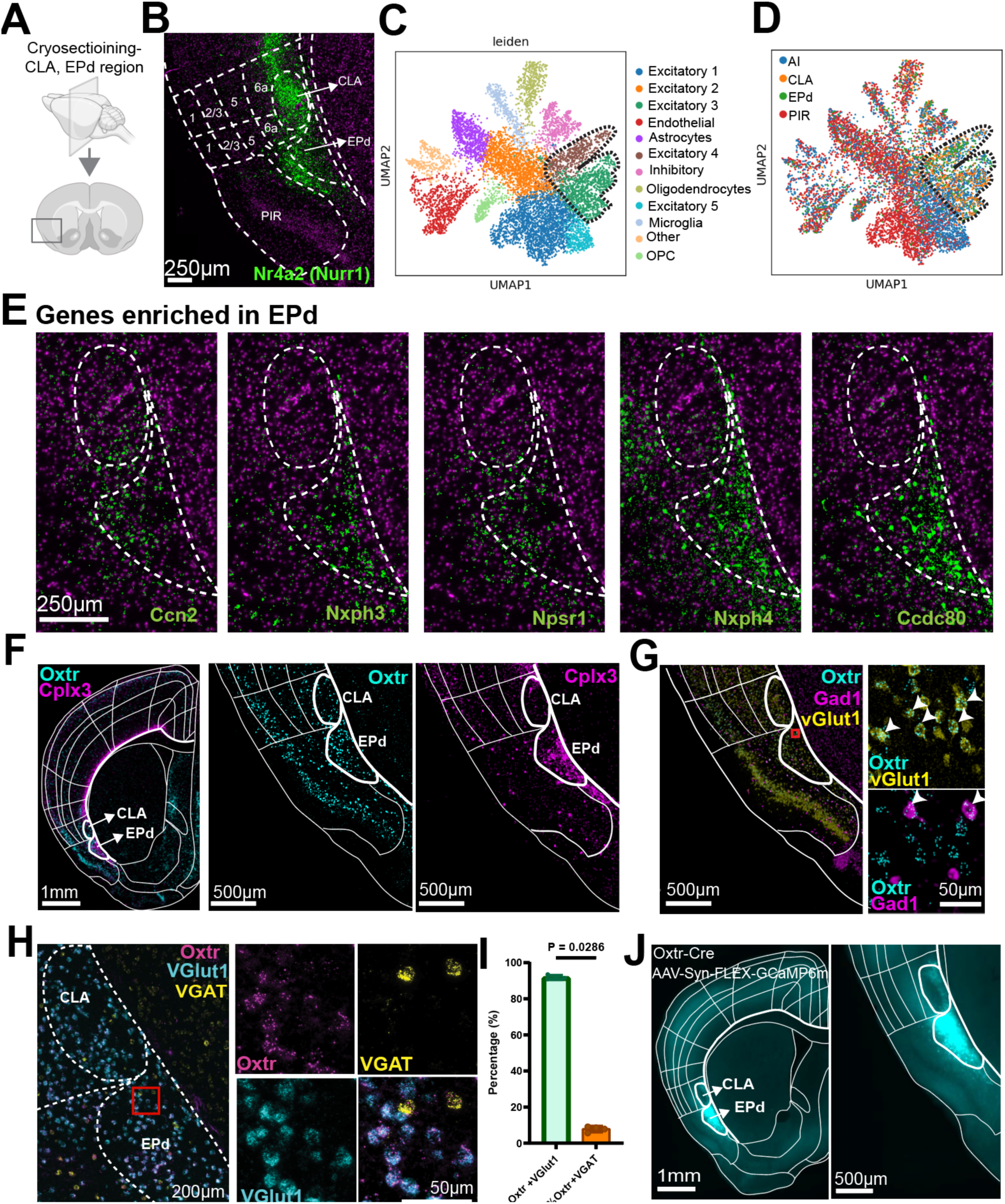
Molecular signature of the EPd with Oxtr enrichment in the mouse brain. **A**. Coronal cryosectioning from the anterior EPd (bregma: +0.5) for spatial transcriptomics. **B**. Enrichment of Nr4a2 (Nurr1), a cortical subplate marker, with elevated expression in the CLA and EPd. **C**. UMAP clustering of all cells from the agranular insular area (AI), CLA, EPd, and piriform area (PIR). **D**. Cells from AI, CLA, EPd, and PIR overlaid on the UMAP. The dotted line highlights excitatory clusters 2 and 3, primarily composed of CLA and EPd. **E.** Genes enriched in the EPd compared to the CLA, revealed through spatial transcriptomics of the same coronal section. **F**. Oxtr is enriched in the EPd, co-expressed with another EPd marker, Cplx3. **G**. Colocalization of Oxtr with Gad1 and vGlut1 using spatial transcriptomics, showing that the majority of Oxtr neurons are excitatory (vGlut1 positive). **H**. Independent validation by RNA in situ hybridization, demonstrating colocalization of Oxtr, vGlut1, and VGAT. **I**. Quantification of the colocalization of Oxtr-positive neurons with vGlut1 and VGAT (n = 4 mice, 3 sections each). P-values were obtained using the Mann-Whitney test. **J**. Oxtr-Cre mice were injected with AAV-Syn-FLEX-GCaMP6m, resulting in localized expression in the EPd.

### Oxytocin receptor positive neurons are enriched in the EPd

Lack of clear molecular markers to label EPd neurons presents a major challenge to understand anatomical and functional organization of the EPd. In our previous Oxtr mapping in the whole mouse brain, we coincidentally identified enriched expression of Oxtr (+) neurons in the EPd(*18*). To ascertain the previous finding, we conducted spatial transcriptomics experiments employing Oxtr and other genes associated with anatomical markers, with a focus on the anterior CLA-EPd complex (∼bregma: +0.7mm) in adult C57BL/6J mice. We performed precise image registration to align the Allen Common Coordinate Framework (CCF) labels to an individual section using striatal and cortical layer-specific markers (Supplementary Figure1D). We confirmed the robust expression of Oxtr, primarily in the EPd that was also marked by high expression of the EPd marker, Cplx3(*29*) (Figure 1F). We also identified that Oxtr neurons in the EPd are mostly glutamatergic (vGlut1 positive) (Figure 1G). To quantify this, we employed RNA in situ hybridization (RNA scope) (Figure 1H). We found that a significant majority (approximately 90%) of Oxtr neurons are excitatory characterized with vesicular glutamate 1 (vGlut1) expression while the remaining 10% are inhibitory neurons with vesicular GABA transporter (vGAT) expression (Figure 1I), conforming to the spatial transcriptome data (Figure 1G) and a previous report (*32*).

Next, we tested whether Oxtr-Cre mice can be used as a transgenic animal tool to selectively label EPd neurons when combined with stereotaxic injection of conditional reporter virus (AAV-Syn-FLEX-GCaMP6m) into the EPd area (Figure 1J). Indeed, viral reporter gene expression is confined in the EPd, hence serving as a reliable tool to understand neuroanatomical and functional organization of the area.

### EPd neurons mainly project to the ventral half of the brain

Elucidating long-range output connectivity can help infer anatomical areas under the control of the EPd for various functional roles(*33*, *34*). Hence, we sought to create a detailed anterograde projection map originating from EPd neurons labeled by Oxtr-Cre mice and analyze their functional significance.

We injected a Cre-dependent adeno-associated virus (AAV2-CAG-Flex-eGFP), covering different anterior-posterior areas of the EPd (N=10 animals; one injection per animal; Figure 2A-B). We used serial two-photon tomography (STPT) to image the whole brain at single cell resolution(*19*, *35*). We detected projection signals using machine learning algorithms and registered individual samples onto the Allen CCF (Figure 2C-D; Supplementary Movie 1). Lastly, maximum projection data from all samples were used to represent the efferent output from the EPd (Figure 2D-E; Supplementary Movie 1). Our analysis revealed abundant projections from EPd neurons to the ventral half of the telencephalon, with sparse projections to the diencephalon and no projection to the midbrain and the hindbrain (Figure 2C-E). We grouped the areas with long-range projections based on their known function to understand downstream circuits under the control of the EPd.

**Figure 2.**
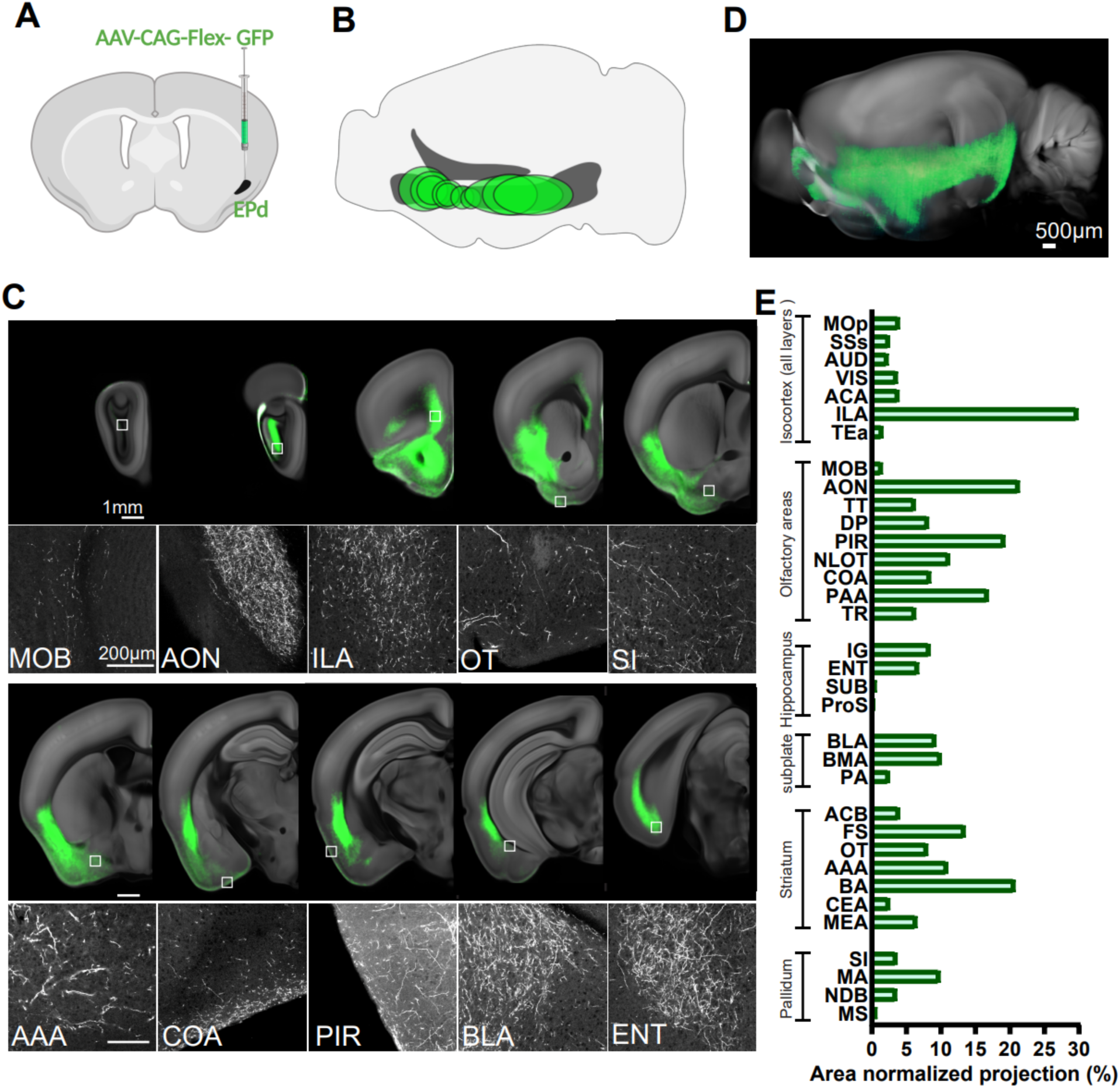
EPd neurons primarily project to the ventral half of the brain. **A**. Schematic of AAV-CAG-Flex-GFP injections into the EPd of Oxtr-Cre mice. **B**. Locations of injections along the anterior-posterior axis of the EPd and the size of individual injections. **C**. Examples of long-range projections (green) from EPd-Oxtr neurons. The bottom panel shows high-magnification images of the white-boxed areas in the top panel. **D**. Maximum projection outputs (from all injections, n=9 mice) from the EPd, registered to the Allen CCF. **E**. Bar graph representing major output areas throughout the brain. The area-normalized projection represents the ratio of the area covered by the projection signal to the total area of the region of interest (ROI), shown as a percentage. Full names of abbreviations according to AllenCCF are listed in Supplementary Table 1.

The major projection areas were olfactory regions, particularly the main olfactory epithelium pathway to process volatile odor cues including the main olfactory bulb (MOB) and its direct downstream areas (e.g., the anterior olfactory nucleus; AON, the piriform area; PIR, and the olfactory tubercle; OT)(*36*). In contrast, the vomeronasal pathway to process mechanical cues (e.g., pheromone) received no projection (e.g., the accessory olfactory bulb; AOB) or sparse projection (e.g., cortical amygdala posteriormedial; COApm) from the EPd(*37*). Another area with prominent projection is the basal forebrain area (magnocellular area; MA, substantia innominata; SI, nucleus of the diagonal band; NDB, and medial septum; MS), which has been implicated in attention and reward processing(*38*). Moreover, significant projections were observed in the limbic areas, including the central amygdala (CEA), basolateral amygdala (BLA), medial amygdala (MEA), bed nucleus of stria terminalis (BNST), infralimbic cortex (ILA), and ventral subiculum (SUBv) that regulate affective behavior(*39*). In contrast, the majority of isocortical and dorsal hippocampal areas that regulate cognitive behavior receive very sparse or no projection. These output connectivity patterns suggest that EPd neurons may play a significant role in regulating olfactory and limbic information processing.

### EPd neurons project selectively to deep layers of the lateral association and medial prefrontal cortices

Previous projection mapping of CLA has revealed abundant isocortical projections to all layers(*5*). To compare how EPd projections differ from CLA projections, we analyzed layer-specific projection patterns to all the isocortical areas (Figure 3). Our analysis revealed that the majority of isocortical areas receive sparse projections, except the lateral association areas (agranular insular; AI, gustatory; GU, perirhinal; PERI, visceral; VISC, and ectorhinal areas; ECT) and the medial prefrontal cortex (prelimbic; PL, infralimbic; ILA, orbital; ORB) (Figure 3A-C). Notably, the EPd sends little to no projection to the retrosplenial cortex (RSP) where the CLA heavily projects to(*5*) (Figure 3C). Additionally, the EPd only projects to deep layers, predominantly layer 6, of the target areas (Figure 3C-D), which contrasts sharply with the CLA that projects to all layers of the isocortex(*5*). Layer 6 (L6) contains neurons with cortical thalamic projection(*40*, *41*). Hence, selective L6 innervation by EPd neurons can regulate corticothalamic output largely from limbic cortical areas(*42*).

**Figure 3.**
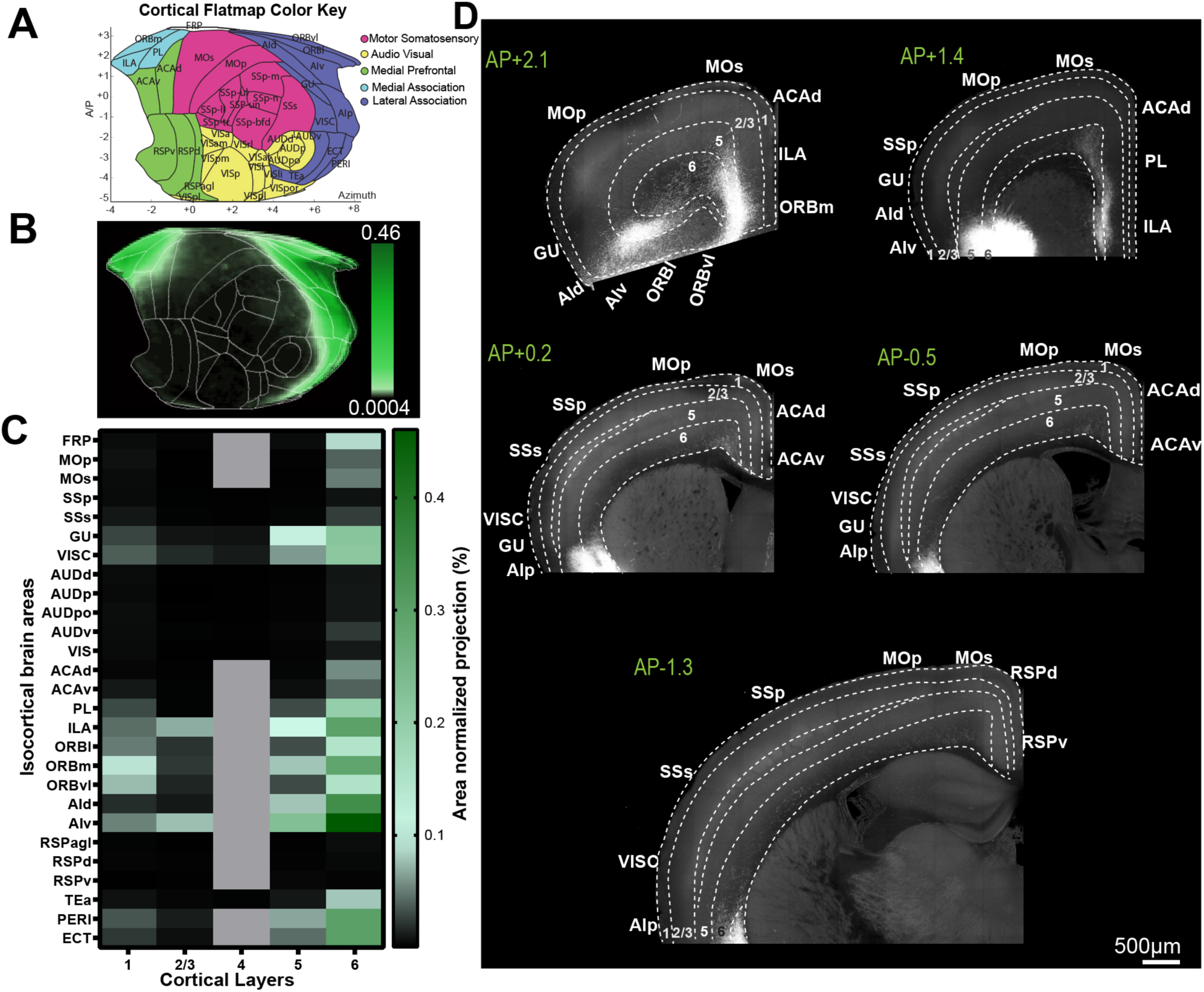
EPd neurons selectively project to the deep layers of the lateral association and medial prefrontal cortices. **A.** Isocortical flatmap showing Allen CCFv3 anatomical regions and border lines. **B**. Isocortical flatmap depicting the projection density from the EPd. **C**. Heatmap of area-normalized projections (percentage of area with signal/total area) from the EPd to isocortical layers. Full names of abbreviations according to AllenCCF are listed in Supplementary Table 1. **D**. Examples of raw images showing long-range projections (green) from the EPd to the isocortex.

### The EPd is topographically organized to project to downstream areas

The EPd can broadly project to downstream areas regardless of their location along the anterior-posterior axis or subregions of the EPd may have preferential projections to a subset of downstream areas. To further understand anatomical organization of the EPd, we crossed Oxtr-Cre mice with Ai65 mice (Cre and Flp dependent intersectional reporter)(*43*) and administered rAAV-EF1a-DIO-FLP-WPRE-hGH polyA injections into one of four major projection areas of the EPd: two anterior brain regions (anterior olfactory nucleus; AON, ILA) and two posterior brain regions (cortical amygdala; COA, Piriform area; PIR) (Figure 4A). We used serial two-photon tomography to examine labeled cells in the whole brain (Figure 4B). Subsequent analysis revealed that injections into AON and ILA predominantly labeled neurons in the anterior parts of the EPd, while injections into COA and PIR resulted in labeling of neurons in the posterior parts (Figure 4B-C), indicating topologically distinct projection domains within the EPd. We also found that both the CLA and the EPd project to the AON and the ILA (Figure 4B-C). In contrast, the PIR and the COA received input predominantly from the EPd (Figure 4B-C).

**Figure 4.**
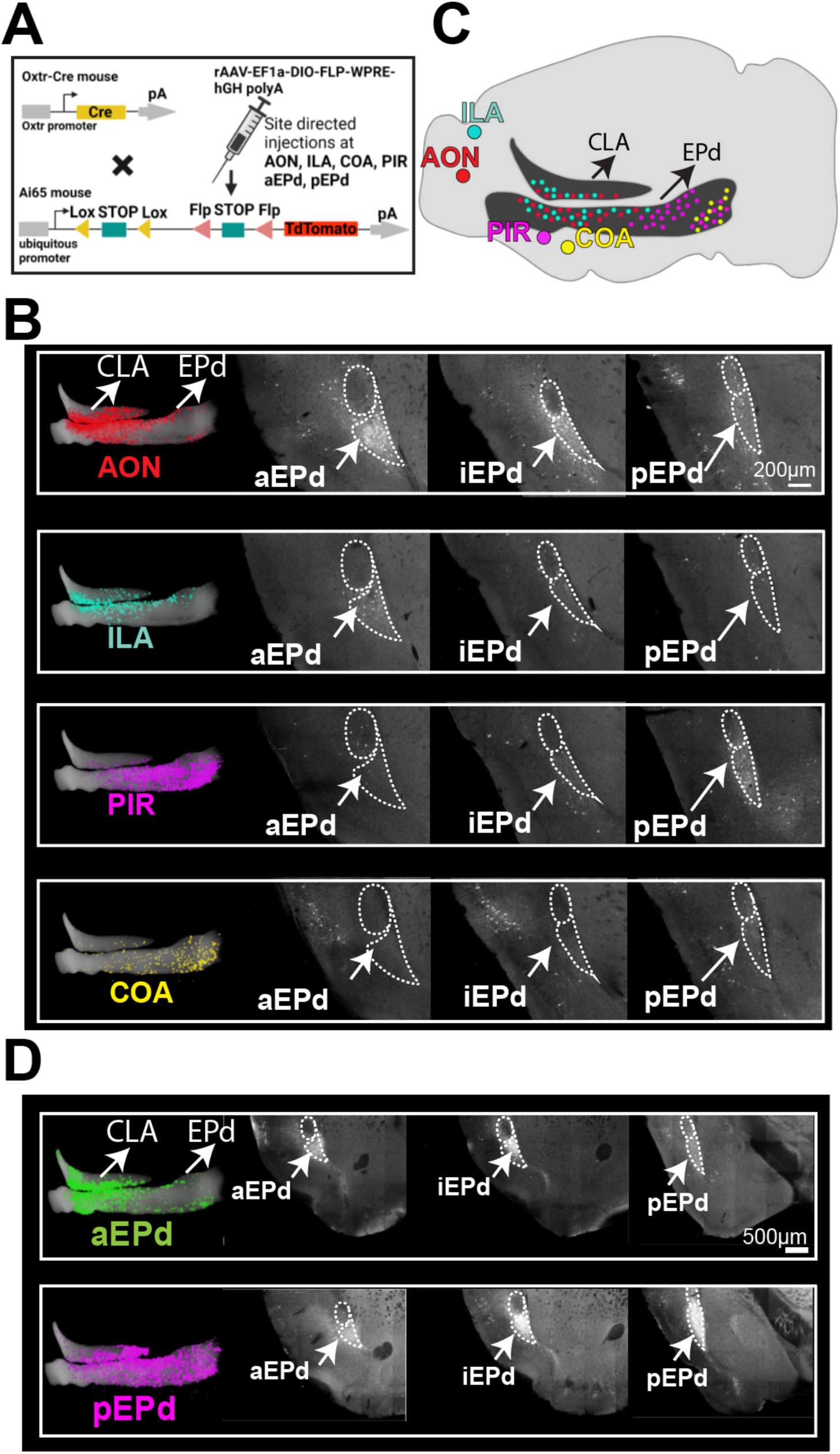
The EPd is topographically organized to project to downstream areas. **A**. Schematic illustrating the breeding and injection strategy. **B**. Examples of raw images showing Oxtr-positive neurons across the anterior to posterior extent of the CLA and EPd, projecting to the AON, ILA, PIR, and COA. **C**. Summary figure depicting CLA and EPd Oxtr neurons projecting to the AON, ILA, PIR, and COA. Larger color-filled circles indicate the injection area, while smaller circles of the same color represent Oxtr-positive cells in the CLA and EPd that project to the respective areas. **D**. Topographic organization of projections within the EPd. Anterior EPd injections show stronger projections in the anterior regions of the CLA and EPd, whereas posterior injections show more projections within the posterior regions of the CLA and EPd. Abbreviations: aEPd: anterior Epd, iEPd: intermediate EPd and pEPd: posterior EPd.

CLA neurons are known to have extensive inter-connectivity rostrocaudally within the structure(*44*). Hence, we hypothesized that EPd neurons will also exhibit uniform rostrocaudal connectivity within the structure. To test this, we injected rAAV-EF1a-DIO-FLP-WPRE-hGH polyA into either the anterior or posterior EPd of the Oxtr-Cre mice, followed by STPT imaging. The results revealed that the anterior EPd (aEPd) exhibited stronger connections to anterior regions of the complex, whereas the posterior EPd (pEPd) injections showed stronger connections to posterior regions (Figure 4D). Altogether, these findings demonstrate an anatomical topology in the connections of the EPd along its rostro-caudal axis, as well as their projections to downstream areas.

### The EPd receives monosynaptic inputs mainly from the limbic and olfactory processing areas

To understand the immediate upstream inputs of the EPd, we performed brain-wide monosynaptic input tracing using conditional AAV expressing TVA and optimized G protein with mCherry reporter injected into the EPd of Oxtr-Cre mice, followed by G deficient avian rabies viruses with GFP expression in the same area (Figure 5A)(*19*) (see Methods for more details). Ten mice were used, with different injection sites, covering the entire anteroposterior extent of the EPd (Figure 5B). We used tissue clearing that preserves endogenous fluorescent protein(*45*), followed by light sheet fluorescence microscopy (LSFM) imaging with cellular resolution. We utilized our automated cell counting methods to quantify fluorescently labeled cells and registered individual samples onto the Allen CCF (Figure 5C). We confirmed specific input cell labeling in the EPd and samples with ∼95% starter cells in the EPd were only included in the analysis (Figure 5D).

**Figure 5.**
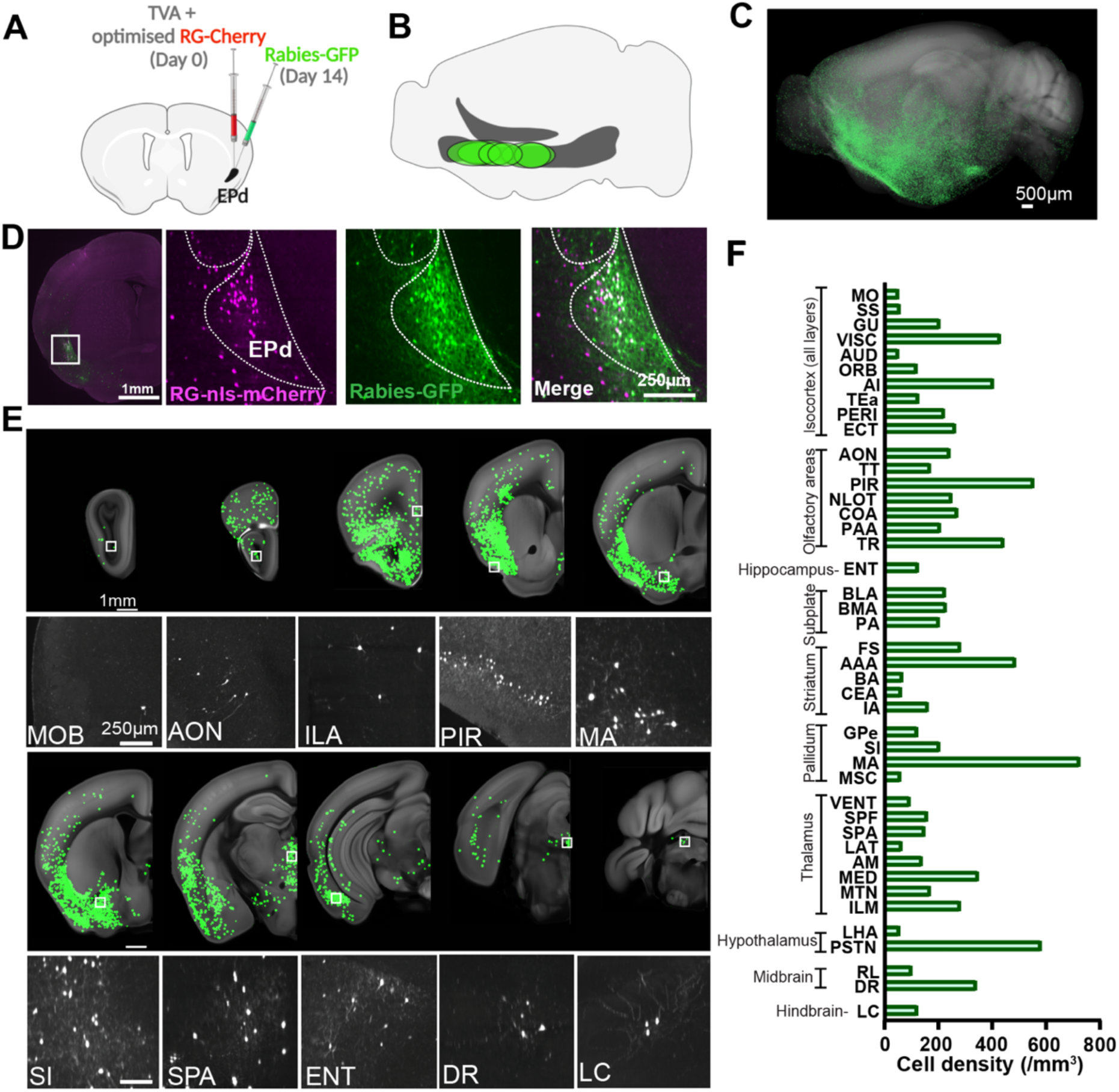
The EPd receives monosynaptic inputs primarily from limbic and olfactory processing areas. **A.** Schematic of conditional monosynaptic tracing using rabies virus injections into the EPd of Oxtr-Cre mice. **B**. Locations of injections along the anterior-posterior axis of the EPd and the size of individual injections. **C**. Brain-wide inputs into the EPd (green, n = 10 animals) targeting Oxtr neurons. Maximum signals from all samples were overlaid on the reference brain. **D**. Starter cells showing colocalization of nls-mCherry and Rabies-GFP in the EPd. **E**. Representative monosynaptic inputs in different coronal planes (top panel) with high-magnification images from the white-boxed areas (bottom panel). **F**. Bar graph representing major brain regions providing monosynaptic inputs to the EPd. Full names of abbreviations according to AllenCCF are listed in Supplementary Table 1.

Overall, EPd neurons receive strong input from the limbic and olfactory areas, similar to the projection mapping, indicating very strong reciprocal connectivity (Figure 5E-F; Supplementary Movie 2). In addition, we found strong input from many thalamic areas such as the medial dorsal thalamus (MED) and the intralaminar thalamus (ILM), linked with limbic, pain, emotional, and autonomic functions (Figure 5E-F; Supplementary Movie 2)(*46–50*). Notably, the EPd receives strong input from basal forebrain, particularly, the magnocellular area (MA) that contains cholinergic neurons broadly projecting to the olfactory areas and ventral hippocampus(*51–53*). In the hypothalamus, the parasubthalamic nucleus (PSTN) that regulates emotion and autonomic functions(*54*, *55*) provides strong input to the EPd. In the mid-brain, the substantia nigra reticular (SNr) dorsal part and the superior collicular (SC) lateral part that receive input from limbic cortices(*56*, *57*), and the dorsal raphe (DR) for emotional control(*58*, *59*) provide input to the EPd (Figure 5E-F; Supplementary Movie 2). In the hindbrain, the locus coeruleus (LC), which provides norephinephrine projections for attention(*60*, *61*), sends input to the EPd (Fig 5E, F; Supplementary Movie 2).

Lastly, we closely examine monosynaptic input from the isocortical areas using our isocortical flatmap (Figure 6A-B). Overall, there were relatively sparse inputs from the isocortex, with the highest inputs from the deep layers (layers 5 and 6) of lateral association areas (Figure 6A-D). This suggests that the EPd mainly receives limbic and internal (gustatory; GU, Visceral cortex; VISC) information from the isocortex.

**Figure 6.**
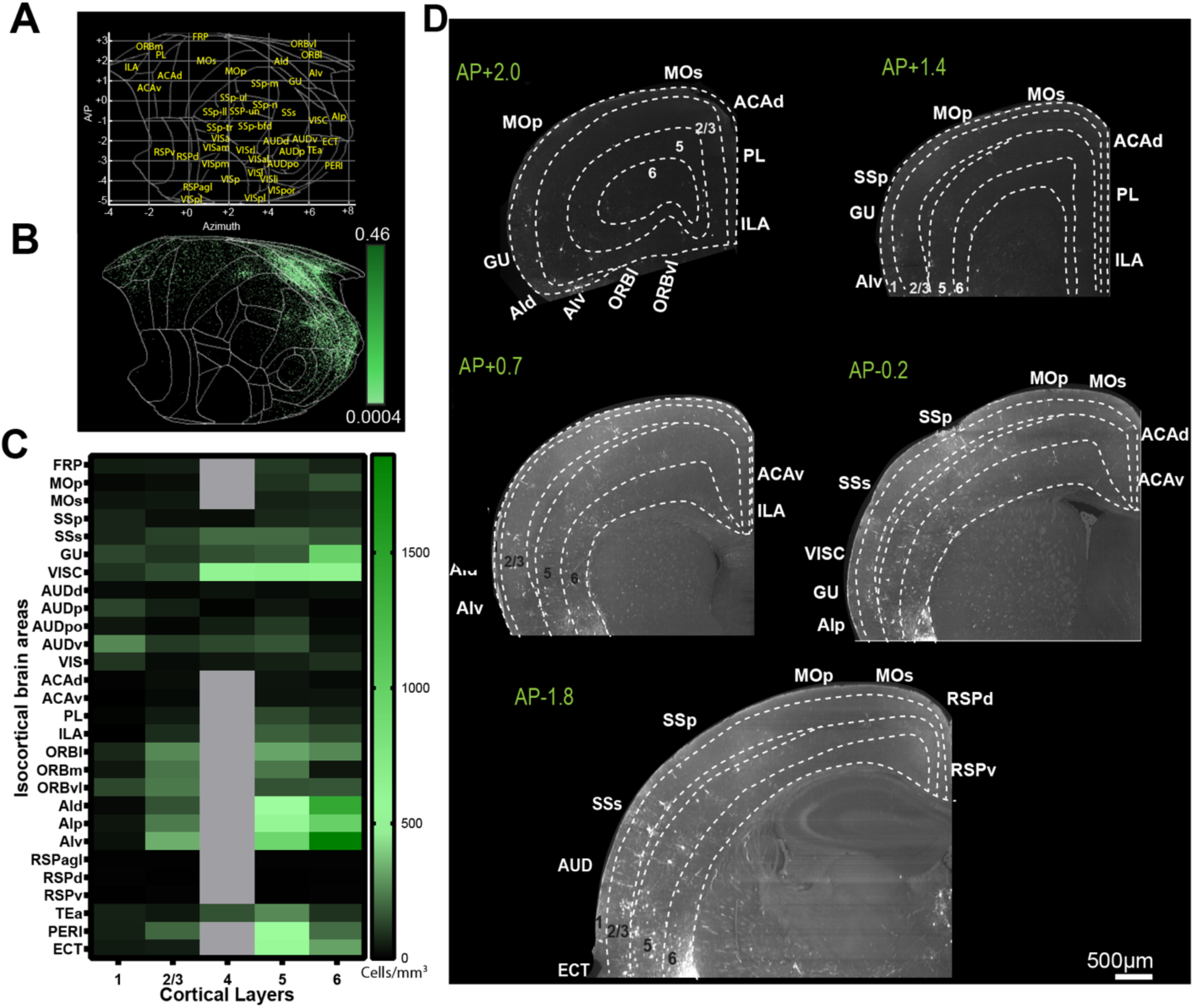
The EPd receives monosynaptic inputs selectively from the limbic and lateral cortices. **A**. Isocortical flatmap showing Allen CCFv3 anatomical regions and border lines. **B**. Isocortical flatmap depicting monosynaptic inputs to the EPd from the isocortex. **C**. Heatmap showing the density of monosynaptic input cells from isocortical layers to the EPd. Full names of abbreviations are listed in Supplementary Table 1. **D**. Examples of raw images showing monosynaptic inputs from the isocortex to the EPd.

Collectively, this data suggests that the EPd receives monosynaptic input from brain areas processing internal state including interoception.

### EPd neurons show downregulated activity upon novel cue exposure

The EPd is a part of the claustrum complex that has been previously implicated in salience processing(*62*). Monosynpatic input data also showed strong input from the salience network (e.g., amygdala, anterior insular)(*63*, *64*). Moreover, it is well established that oxytocin signaling has been implicated in regulating environmental novelty for both social and non-social stimuli(*65*– *68*). Based on this evidence, we hypothesized that Oxtr-positive EPd neurons will be activated upon social cues.

To examine the neural activity patterns of the EPd neurons to social cues, we injected AAV-Syn-FLEX-GCaMP6m into the anterior EPd and implanted a graded index (GRIN) lens with a miniscope (Figure 7A; See Methods for more details). We exposed our target mice to a novel social cue (an opposite-sex stranger mouse) in a home cage while recording GCaMP6m signals (Figure 7B-E). In contrary to our original prediction, we observed high baseline neural activity during exploratory behaviors in home cage followed by significantly decreased neural activity upon encountering a stranger mouse (Figure 7B-E). The neural activity returned to high baseline levels in about 50 sec after, which was maintained during subsequent no-stimulus periods (Figure 7C-D). Further, to determine if this decrease in neuronal activity was dependent on social or non-social novelty, we introduced a novel object. Similar to the social novelty cue, there was a significant decrease in activity upon introducing a novel object (Figure 7D-E).

**Figure 7.**
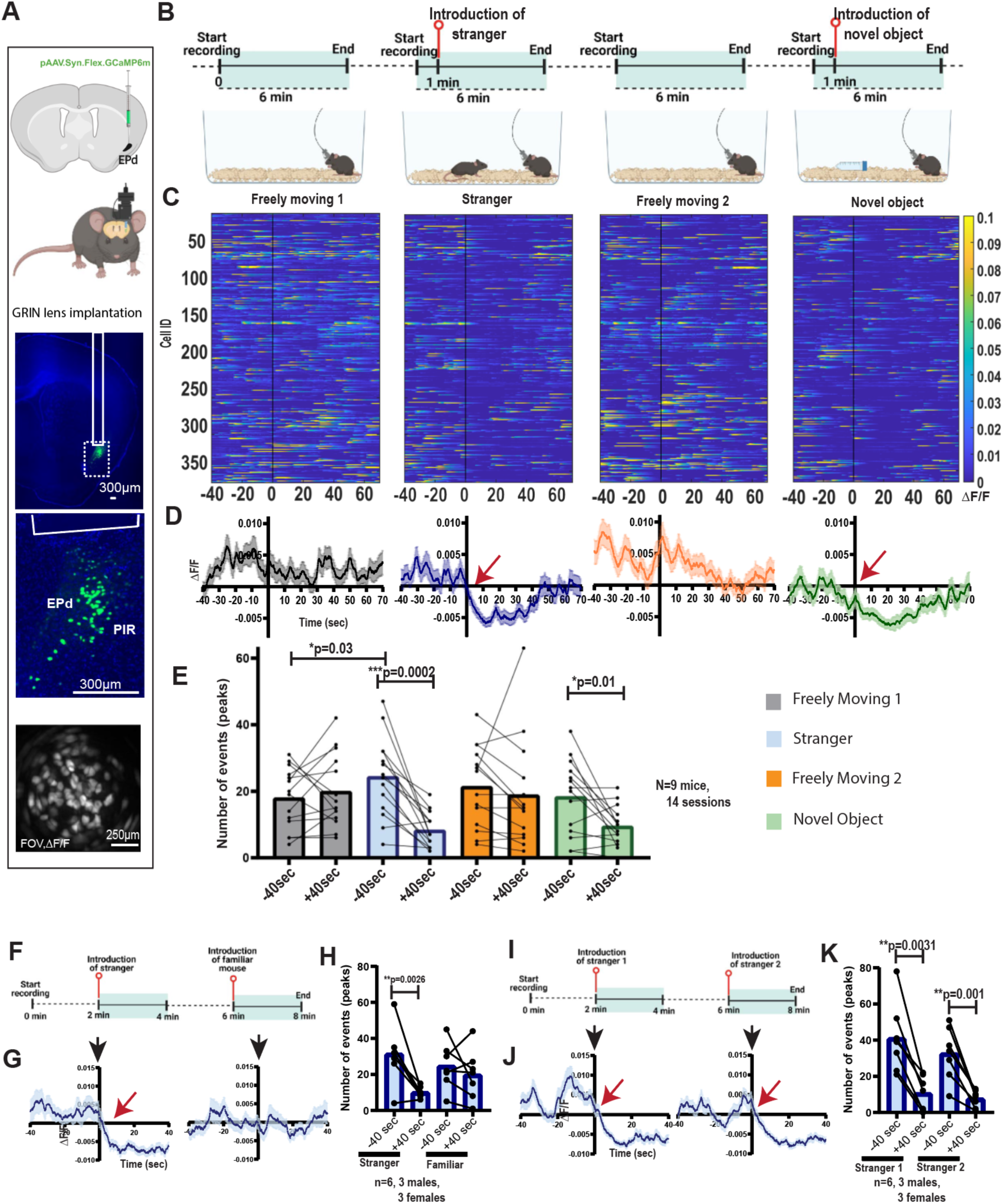
EPd neurons exhibit downregulated activity upon exposure to novel cues. **A**. Top: GCaMP6m virus injection into the EPd of Oxtr-Cre mice. Bottom: GRIN lens implantation in the target area and an example of in vivo imaging. **B**. Behavioral paradigm. **C**. Heatmap showing all neurons recorded 40 seconds before and 70 seconds after the event. **D**. High and sustained neuronal activity (ΔF/F) in the home cage (without external cues) and significantly decreased activity upon exposure to novel cues. **E**. Number of calcium events before and after 40 seconds of the event, analyzed using the Wilcoxon matched-pairs signed-rank test. **F-G**. Repeated exposure to the same stranger mouse (F) did not cause a significant decrease in EPd neural activity (ΔF/F). **H**. Number of calcium events before and after 40 seconds of introducing either a stranger or familiar mouse, Wilcoxon matched-pairs signed-rank test. **I-J**. In contrast, introducing two different stranger mice (I) led to continued downregulation of Oxtr neuronal activity (ΔF/F). **K**. Number of calcium events before and after 40 seconds of introducing two stranger mice, Wilcoxon matched-pairs signed-rank test.

To test whether novelty drives the down-regulation of EPd activity, we repeatedly exposed the mice to the same stranger mouse (Figure 7F-H) or to two different stranger mice (Figure 7I-K). Repeated exposure to the same mouse abolished the significant decrease in EPd activity. In contrast, exposure to a new stranger mouse again significantly reduced activity, confirming that the novelty of the social stimuli causes the down-regulation of the EPd activity (Figure 7I-K). Similar to the results from repeated social cue introductions, repeated exposure to the non-social cue (toy mouse) abolished the downregulation of neuronal activity (Supplementary Figure 2A-D), indicating that novel stimuli (social or non-social) downregulated activity of Oxtr-positive EPd neurons.

## Discussions

We present a comprehensive cellular characterization and anatomical connectivity map of the EPd, a highly enigmatic brain area. We found strong connections from the EPd to the ventral half of the brain including limbic and olfactory areas. Our in vivo recording shows selective down-regulation of EPd neuronal activity upon novel stimuli from high baseline activity, suggesting its potential role in mediating interoceptive states, which is attenuated by external cues. Collectively, our study provides an initial anatomical and functional characterization of the EPd to elucidate its role in the brain.

In rodents, the CLA and the EPd are identified as separate structures, whereas in primates, they appear as a single continuous structure(*69*). This has led to ongoing debates over the anatomical boundaries of the CLA complex in rodents as well as inconsistent nomenclature(*21*, *70*), complicating the selection of a specific marker exclusive to EPd. Using spatial transcriptomics and fluorescence in situ hybridization, we identified distinct gene expression signatures of the EPd and achieved clear delineation of anatomical boundaries between the EPd and the adjacent structures (the CLA, AI and PIR). Notably, we found Oxtr as an additional marker for the EPd in mouse brains(*18*, *32*). Previously, Oxtr expression has been reported in the CLA complex across species, including rodents(*32*, *71*) and primates such as rhesus macaques(*72*). Moreover, our current study identified that EPd-Oxtr neurons are largely glutamatergic neurons, consistent with the previous report(*32*).

We found that Oxtr-Cre mice can serve as a genetic tool to selectively target EPd neurons. Combined with viral tools and 3D mapping methods, we established brain-wide input-output connectivity maps of the EPd. While the input-output architecture of the CLA(*5*, *21*) is widely studied, there is a lack of comprehensive studies that explain the connectivity of the EPd across the whole mouse brain(*73*). Both the CLA and the EPd originate from the same part of the pallium based on Nr4a2 expression during development, with neurons that migrate to the ventrolateral dorsal pallium forming the CLA and that migrate further ventrally forming the EPd(*3*). Due to their common origin, these areas were thought to have a similar topographical organization(*1*). However, recent studies suggest regional differences in the input-output architecture of the CLA complex, with CLA exhibiting a dorsoventral organization in its output projections(*74*). In this study, we identified extensive projection of the EPd neurons in the ventral half of the brain broadly covering the olfactory and limbic areas, which contrasts strongly with the CLA output covering the dorsal half of the brain including the entire isocortical areas (*5*, *21*). Moreover, our projection mapping from the EPd neurons shows no projections to the retrosplenial area (RSP), which is a defining feature of CLA output(*75*). For the monosynaptic input, the EPd receives strong input from limbic and olfactory areas, suggesting a reciprocal connection between the areas. In contrast, thalamic, midbrain, and hindbrain areas that regulate attentive state (MED, LC) and process emotion/affective/pain/visceral information (subparafascicular nucleus, PSTN, DR) provide monosynaptic input without receiving reciprocal projection from the EPd. Another noticeable area is the basal forebrain (e.g., SI, MA) which contains cholinergic neurons to mediate attention(*38*, *76*). Collectively, anatomical connectivity data strongly suggest that the EPd is positioned to modulate limbic and olfactory areas based on input from emotional and visceral circuits, serving as a parallel circuit to the CLA for motor sensory processing in response to external stimuli(*2*).

Previous studies suggest that the EPd is implicated in recognition memory(*77*), epilepsy(*13*, *78*) and higher order olfactory processing(*79*). Our *in vivo* recording data showed a high level of neural activity at the baseline exploratory stage and persistent down-regulation of the EPd activity upon novel stimuli (both social and non-social). Hence, our data suggest that the EPd may regulate baseline exploratory behavior while inhibition of the EPd allows attentive neural circuits to engage in response to novel stimuli. Previous studies showed that long-range projections from the CLA excitatory neurons preferentially target local interneurons in cortical areas, resulting in net inhibition of the downstream circuit(*80*, *81*). Moreover, inhibition of the CLA impaired performance selectively in high order decision making(*82*, *83*). Similarly, the EPd may provide top-down inhibition to the olfactory and limbic circuit, and inhibition of the EPd can disinhibit the downstream circuit to express initial cautious and anxious approach behavior upon novel stimuli while considering internal state information. Another interesting observation is high baseline neural activity of the EPd, which suggests the EPd’s involvement in the interoceptive state. Indeed, a prior study identified very high level of EPd neural activity during slow wave sleep (or deep sleep) compared to awake or REM sleep(*84*). This evidence suggests that the EPd may actively participate in maintaining the baseline interoceptive state, and potentially even deep sleep, serving in opposition to neural circuits that mediate exteroception. The ability to shift between interoceptive and exteroceptive states is vital for assessing internal states and determining appropriate responses to potential environmental threats or opportunities. Impairment of such function can result in pathological conditions such as anxiety or hypervigilance. Indeed, neuroimaging data from patients with schizophrenia show abnormal activity within the CLA complex, which corresponds to the EPd(*85*).

Our study is not without limitations. The EPd contains additional cell types beyond Oxtr+ neurons, each with distinct local and long-range connectivity. Investigating their synaptic connections and elucidating their functional roles will enhance our understanding of EPd circuits. Furthermore, exploring how Oxt signaling influences EPd circuits and how the EPd is altered in various pathological conditions will be critical for future research. We envision that our anatomical and functional studies provide a foundation for these future investigations.

## Methods

### Animals

All animal care and experimental procedures were approved by the Penn State University Institutional Animal Care and Use Committee (IACUC). The Oxtr-Cre line, originally established by Hidema et al. (*86*), was imported to Penn State University via mouse rederivation. Oxtr-Cre mice, which have a 129 × C57BL/6J mixed genetic background, are not commercially available. C57BL/6J mice were utilized for RNA in situ hybridization and spatial transcriptomics analysis. Mice were provided food and water ad libitum and were housed under controlled temperature and light conditions, with a 12/12-hour light/dark cycle.

### Spatial Transcriptomic Analysis

#### Sample Collection

C57BL/6J mice, aged 3-4 months, were euthanized via cervical dislocation to collect fresh frozen whole brain samples for MERFISH experiments. After excision, brains were placed in cryomolds (Ted-Pella, 27110) partially filled with Optimal Cutting Temperature (O.C.T.) compound (Tissue-Tek 4853). The molds were then filled completely and rapidly frozen using 2- methylbutane cooled by dry ice. Once the O.C.T. compound had solidified, the samples were stored at –80°C for long-term preservation. Dissection tools were disinfected with RNaseZap (Invitrogen, AM9780) before and after each use to ensure RNA integrity.

#### MERFISH 500 Gene Panel Selection

A 500-gene panel was selected for the MERFISH experiments. This panel included cell-type markers and genes specifically enriched in regions such as the claustrum (CLA), dorsal endopiriform nucleus (EPd), piriform cortex (PIR) and agranular insular cortex (AI). The panel also comprised cortical layer-specific markers (refer to Supplementary Table 2 for details).

#### MERFISH Imaging Experiment

MERFISH experiments were performed using the Vizgen MERSCOPE platform. Fresh frozen mouse brain samples were cryosectioned at a thickness of 10 μm using a cryostat maintained at −20°C. Sections were mounted on MERSCOPE slides (Vizgen 2040000) and fixed in 4% paraformaldehyde (PFA) in 1x PBS for 15 minutes. Following fixation, sections were washed three times with 1x PBS and permeabilized in 70% ethanol at 4°C overnight. Probe hybridization was carried out with a custom 500-gene MERSCOPE Gene Panel Mix (Vizgen 20300008) in a 37°C incubator for 36-48 hours. Post-hybridization, sections underwent two 30-minute washes with Formamide Wash Buffer at 47°C. Samples were then embedded in a hydrogel using the Gel Embedding Premix (Vizgen 20300004), ammonium persulfate (Sigma 09913-100G), and TEMED (Sigma T7024-25ML) from the MERSCOPE Sample Prep Kit (10400012). Once the hydrogel solidified (∼1.5 hours), the samples were cleared overnight at 37°C with a Clearing Solution containing Proteinase K (NEB P8107S) and Clearing Premix (Vizgen 20300003). After clearing, sections were stained with DAPI and Poly T Reagent (Vizgen 20300021) for 15 minutes at room temperature, followed by a 10-minute wash in Formamide Wash Buffer. Imaging was then performed using the MERSCOPE system (Vizgen 10000001). Comprehensive protocols for sample preparation and instrument usage can be found at Vizgen Resources and Vizgen Instrumentation.

#### Registration to Allen CCF

The Allen Common Coordinate Framework (CCF)(*87*) was used to register all brain slices that underwent MERFISH experiments. Initially, datasets were loaded individually into the MERFISH Visualizer, where images with DAPI and other markers were exported, along with a metadata file detailing image size and center coordinates in microns. Each brain slice was then aligned to the correct plane of the CCF in all dimensions using ITK-Snap, with manual adjustments to the reference brain’s orientation. Subsequently, registration was performed using Applied Normalization Tools (ANTs)(*88*), which applied a combination of Rigid, Affine, and Deformable SyN transformations to achieve precise alignment. For final refinements, ImageJ’s BigWarp plugin(*89*) was employed to ensure optimal registration accuracy. A label image was created to delineate brain regions, with each pixel assigned a corresponding region value. Custom MATLAB code was then used to convert the x and y micron coordinates of each cell into pixel coordinates, allowing cells to be assigned to specific brain regions based on the corresponding CCF label image.

#### Preprocessing

Preprocessing was performed using the Scanpy framework in Python(*31*). Data from three replicates were pooled, and the gene set was narrowed down to 247 genes related to cerebrovascular function, claustrum and dorsal endopiriform nucleus marker genes, neurotransmitter receptors/transporters, cell type markers, energy-related genes, and cortical markers. Cells from the claustrum, dorsal endopiriform nucleus, agranular insula, or piriform areas were filtered. Doublets were identified and removed using Scrublet(*90*), and cells smaller than 40 µm or larger than three times the median size were excluded. Additional filtering removed cells expressing fewer than 5 genes, fewer than 10 total transcripts, and genes expressed in fewer than 3 cells. Counts were normalized by cell volume, and cells with total RNA counts outside the 2nd and 98th percentiles were removed. To stabilize variance, a logarithmic transformation was applied to the normalized counts. The data were then z-score normalized and values were further rescaled so that the maximum value for each gene was set to 10.

#### Clustering

Clustering was performed by first applying principal component analysis (PCA) for dimensionality reduction. A neighborhood graph was then constructed using 10 nearest neighbors and the first 20 principal components, followed by computing a UMAP embedding to visualize the data in two dimensions. Leiden clustering was performed with a resolution parameter of 0.2 to identify distinct cellular populations. Cell types were subsequently assigned to clusters based on established marker genes: Astrocytes (Aqp4, Slc4a4), Endothelial cells (Ptprb, Vegfc), Microglia (Cx3cr1, Csf1r, Cd68), Oligodendrocytes (Mobp, Plp1, Cldn11), OPCs (Pdgfra, Cspg4), Excitatory neurons (Slc17a6, Slc17a7, Scl17a8), and Inhibitory neurons (Slc32a1, Vip, Sst, Npy, Pvalb).

### Single-molecule mRNA fluorescence in situ hybridization

C57BL/6J mice (2-3 months old) were euthanized by cervical dislocation, and their brains were immediately dissected and immersed in Optimal Cutting Temperature (OCT) media (Tissue-Tek, catalog #4853). The immersed brains were rapidly frozen using dry ice-chilled 2-methylbutane and stored at −80°C until needed. Coronal brain sections, 10 μm thick, were collected using a cryostat and stored at −80°C. In situ hybridization was conducted within one month of sectioning.

The RNA Scope Multiplex Fluorescent Reagent Kit v2 (ACDBio) was employed to detect and quantify target mRNA at single-molecule resolution, following the manufacturer’s protocols for fresh frozen samples. To quantify the colocalization of Oxtr-positive cells with markers for excitatory and inhibitory neurons, Probe-mm-Oxtr (454011-C2), Probe-mm-slc17a7 (ACDBio, 416631), and Probe-mm-VGAT (319191-C3) were used to detect mRNA expression of Oxtr, vGlut1, and vGAT.

### Vibratome sectioning of mouse brain samples

Oxtr Cre mice injected with pAAV.Syn.Flex.GCaMP6m.WPRE.SV40 (AAV1) virus (titer: 2.1 × 10^13 GC/ml), purchased from Addgene and generously provided by Douglas Kim & the GENIE Project (RRID: Addgene_100838) into the anterior part of the EPd (AP: 1.1 mm; ML: 2.45 mm; DV: −4.13 mm) was transcardially perfused with 4% paraformaldehyde (PFA). Brain samples were dissected and stored in 4% PFA overnight. Afterward, the samples were washed four times with 0.05M phosphate-buffered saline, followed by embedding in 4% agarose solution. The embedded samples were sectioned using a vibratome at a thickness of 60 µm. The anterior EPd sections containing the signal were mounted on glass slides with FluorSave reagent and coverslipped. The brain sections were then imaged using a fluorescence microscope.

### Fluorescence microscopy imaging

Microscopic imaging was performed using a BZ-X700 fluorescence microscope (Keyence). A low magnification objective lens (4x) provided a broad view to define brain AP locations from bregma, while a higher magnification objective lens (20x) was used to image the EPd area in each brain section. Images were manually delineated based on the Allen atlas, and colocalization analysis was conducted manually using the Cell Counter plugin in FIJI (ImageJ, NIH).

### Stereotaxic surgery and virus injections

For anterograde tracing, 50–500 nl of AAV2-CAG-Flex-EGFP virus (titer 3.7 × 10¹² vg/ml, purchased from the UNC Vector Core) was injected into the EPd as previously described². Oxtr-Cre mice (2-4 months old, 7 males and 3 females) were anesthetized with isoflurane (administered using Somnosuite, Kent Scientific) and mounted on a stereotaxic instrument (Angle Two, Leica) with a heating pad underneath. All injections were performed with pulled micropipettes (VWR, catalog #53432-706). The virus was delivered through the small opening of the micropipette at a rate of 75–100 nl/min. The speed and volume of injection were monitored using the calibration marks on the micropipette (1 mm = 100 nl). To target the anteroposterior extent of the EPd, two major coordinates were used, and post hoc analysis was performed to confirm the actual injection site. Coordinates for anterior EPd injections were anteroposterior (AP) from the bregma: 1.1 mm; mediolateral (ML): 2.45 mm; dorsoventral (DV): −4.13 mm. For intermediate EPd injections, the coordinates were: AP: −0.7 mm, ML: 3 mm, and DV: −4.3 mm. After 3 weeks, the mice were deeply anesthetized with a ketamine-xylazine mixture (100 mg/kg ketamine, 10 mg/kg xylazine, i.p.) and perfused. Brain samples were then collected by cardiac perfusion with 4% paraformaldehyde (PFA) for serial two-photon imaging.

To study the anatomical topology of EPd neuronal projections, we employed a transgenic approach combined with viral injections. Oxtr-Cre mice were crossed with Ai65 mice (Jax; Stock No: 021875). Adult mice (2-3 months old) were injected with rAAV-EF1alpha-DIO-FLP-WPRE-hGH polyA (BioHippo, catalog #BHV12400173, 200 nl each) into six major target locations: the anterior olfactory nucleus (AON, 2 males and 2 females; coordinates AP: 2.22 mm, ML: 0.7 mm, DV: −4.0 mm), the infralimbic area (ILA, 1 male and 2 females; coordinates AP: 1.70 mm, ML: 0.46 mm, DV: −2.70 mm), the piriform cortex (PIR, 2 females; coordinates AP: −2.56 mm, ML: 3.35 mm, DV: −5.20 mm), the cortical amygdaloid nucleus (COA, 1 male and 1 female; coordinates AP: −1.58 mm, ML: 2.50 mm, DV: −5.61 mm), the anterior EPd (3 males; coordinates AP: 1.10 mm, ML: 2.45 mm, DV: −4.13 mm), and the intermediate EPd (1 male; coordinates AP: −0.7 mm, ML: 3 mm, DV: −4.3 mm). Three weeks post-injection, mice were deeply anesthetized with a ketamine (100 mg/kg) and xylazine (10 mg/kg) mixture and then transcardially perfused with 4% paraformaldehyde (PFA). Brain samples were collected for serial two-photon imaging.

For monosynaptic input tracing, AAV5-CAG(del)>TCIT(-ATG)-Flex(*loxP*)-SV40 (1:8 dilution, titer: 1.6E+12 gc/ml, ref) was co-injected with AAV5-CAG(del)>nC2oG-Flex(loxP)(*91*) (generous gift from Dr. Todd Anthony at Harvard University, titer: 2.8E+12 gc/ml) was injected in Oxtr-Cre mice (*N* = 10 mice-7M and 3F, 2-3 months old) for the whole length of the EPd (150 nl per brain, same co-ordinates as anterograde mapping) as previously described(*19*). After 14d, the same volume of EnvA G-deleted Rabies-EGFP virus (titer: 4.89E+09 TU/ml, purchased from Salk Institute viral vector core, a gift from Edward Callaway; RRID:Addgene_32635; Osakada et al., 2011) was injected into the same location with a 5 degree tilted angle. The mice were euthanized 7 d later for brain collection, tissue clearing and imaging. After all the above-mentioned stereotaxic surgeries, the mice were injected intraperitoneally with 0.2ml of 1mg/ml Carprofen for 3-4 consecutive days to minimize pain.

### STPT imaging and related data analysis

Both anterograde tracing and the anatomic topology of EPd projections were determined using serial two-photon tomography (STPT) imaging and analysis, as previously described(*19*, *92*, *93*). Briefly, dissected brains were postfixed in 4% paraformaldehyde (PFA) overnight at 4°C. The fixed brains were then stored in 0.05 M phosphate buffer (PB) at 4°C until imaging. To image the entire brain, STPT was performed using the TissueCyte 1000 system (TissueVision) following established protocols(*94*, *95*). The brains were embedded in 4% oxidized agarose and cross-linked with a 0.2% sodium borohydride solution. Imaging was conducted with a resolution of 1 × 1 µm² in the x and y dimensions, across 12 × 16 × 280 tiles, with 50-µm intervals in the z-dimension. A wavelength of 910 nm was used for two-photon excitation to simultaneously excite both green (e.g., eGFP) and red (e.g., tdTomato) fluorescent signals. The signals were separated using a 560-nm dichroic mirror and two band-pass filters (607/70-25 for red and 520/35-25 for green). The imaging tiles for each channel were stitched together using custom-built software(*19*, *94*, *96*).

For quantitative projection data analysis, we used our previously published pipeline with minor modifications(*19*). Briefly, both the signal and background channels were z-normalized. The background channel images were then subtracted from the signal channel images to increase the signal-to-noise ratio. Next, we employed Ilastik(*97*) for signal detection. By integrating Ilastik into the automatic workflow, our algorithms performed parallel computations to detect each pixel with the maximum likelihood of it belonging to a projection signal, cell body, brain tissue, or empty space. Projection strength for each area was calculated by summing all projection signal pixels within an anatomically defined region. Autofluorescence of the brains was used to register each brain to the Allen CCF(*87*) using Elastix (*98*), after which the projection signals were transformed to the reference brain. We then used the maximum projection of registered long-range output datasets (N=10) from each area to create a representative projection data set for further quantitative analysis. “Area normalized projection” represents the ratio of the area covered by the projection signal to the total area of the region of interest (ROI), depicted as a percentage. For example, if the total pixel count for one ROI was 20,000 and the area covered by the projection signal for that ROI was 2,000, the area normalized projection would be (2,000/20,000) × 100 = 10%. For EPd anatomical topology distinction, we used the cell body classifier from the Ilastik to count the number of cells, rather than using the projection pixels. We then quantified the cell density per ROI by dividing the total number of cells within one ROI by the total volume of that ROI, as previously described(*92*).

### Tissue Clearing, light sheet imaging and related data analysis

Details of tissue clearing, light sheet imaging, and data analysis are as previously described(*99*). We used these methods for monosynaptic rabies tracing of EPd, primarily employing SHIELD (Stabilization under Harsh conditions via Intramolecular Epoxide Linkages to prevent Degradation) tissue clearing to ensure minimal tissue volume changes while preserving endogenous fluorescence signals(*45*). SHIELD reagents and protocols were obtained from LifeCanvas Technologies (https://lifecanvastech.com/). For P56 brains, PFA-fixed samples were incubated in SHIELD OFF solution for 4 days at 4°C, followed by SHIELD ON buffer for 24 hours at 37°C. Tissues were then incubated in delipidation buffer at 37°C for 10 days, washed overnight in PBS, and optically cleared by sequential incubation in 75% EasyIndex + 25% water for 24 hrs followed by 100% EasyIndex solution at 37°C for 24 hrs. For LSFM imaging, samples were embedded in 2% low-melting agarose (Millipore Sigma, cat. no.: A6013, CAS Number: 9012-36-6) in EasyIndex using a custom holder. They were incubated in EasyIndex at room temperature (20-22°C) for at least 12 hours before imaging with the SmartSPIM light sheet fluorescence microscope (LifeCanvas). During imaging, the holder arm with the sample was immersed in 100% EasyIndex. Our setup included a 3.6X objective lens (LifeCanvas, 0.2 NA, 12 mm working distance, 1.8 μm lateral resolution), lasers at 488 nm, 560 nm, and 642 nm wavelengths, and a 5 μm z-step size. After imaging, samples were stored in 100% EasyIndex at room temperature (20-22°C). We developed a parallelized stitching algorithm for 3D reconstruction, inspired by Wobbly Stitcher(*100*), aimed at conserving hard drive space and minimizing memory usage. The algorithm first captured 10% outer edges of each image tile and generated a maximum intensity projection (MIP) in the z-direction for every set of 32 slices in the stack. It then aligned the z coordinates of MIP images across columns, followed by x and y coordinate alignment. Within each MIP, adjustments to 32 slices were made using curve fitting to finalize tile coordinates.

We enhanced our cell density mapping workflow by integrating the ilastik-based cell detection algorithm with our established methods(*94*, *95*, *101*). Signals smaller than the cellular diameter were filtered out. Centroids of individual cells were then identified. Subsequently, we performed image registration to align cell detection results with the Allen CCFv3(*87*) template using Elastix(*98*). Centroids were counted in each brain region to conduct 3D cell counting. To determine the anatomical volume of each sample, we initially registered the Allen CCF to individual samples with elastix, adjusting anatomical labels accordingly based on registration parameters. We then quantified the number of voxels associated with specific anatomical IDs to estimate the 3D volume of each anatomical area. The density of 3D cell counting per anatomical regional volume (mm³) provided a quantitative measure of cell distribution across brain regions.

### Ca^2+^ imaging with a head-mounted fluorescent microscope

For miniscope-based recordings, adult Oxtr-Cre mice (approximately 9-11 weeks old) underwent two stereotaxic surgeries: one for virus injection and a second for lens implantation. Mice were anesthetized with isoflurane, administered via a Somnosuite (Kent Scientific), and positioned in a stereotaxic frame (Angle Two, Leica) with a heating pad underneath.

The pAAV.Syn.Flex.GCaMP6m.WPRE.SV40 (AAV1) virus (titer: 2.1 × 10^13 GC/ml), purchased from Addgene and generously provided by Douglas Kim & the GENIE Project (RRID: Addgene_100838), was diluted in phosphate-buffered saline (1:3) based on titration experiments to achieve optimal cytoplasmic, non-nuclear localization of GCaMP6m in fixed tissue slices from injected animals. A volume of 200nl was injected into the anterior part of the EPd (AP: 1.1 mm; ML: 2.45 mm; DV: −4.13 mm).

Post-surgery, mice were housed in fresh autoclaved cages with ad libitum food and water. After a recovery period of two weeks, a second surgery was conducted for GRIN lens implantation. An incision was made to expose the skull, and stereotaxic alignment was performed using the inferior cerebral vein and Bregma as vertical references. The same coordinates used for GCaMP injection (AP: 1.1 mm; ML: 2.45 mm; DV: −4.13 mm) were employed for lens implantation. The GRIN lens (Inscopix: ProView™ Integrated Lens, 0.6 mm x 7.3 mm; 1050-004413) was lowered into place at a rate of 200 µm per minute. The lens, attached to a baseplate, was secured to the skull with a small drop of dental cement (C&B Metabond Adhesive Cement system, #S380). After curing, thin layers of dental cement were applied to cover the exposed skull, followed by multiple layers of dental acrylic (Ortho-Jet BCA, Lang Dental) to stabilize the lens and baseplate. Once the dental acrylic had cured, mice were removed from isoflurane and allowed to recover in clean, autoclaved cages to minimize infection risk. Mice with GRIN lens implants were housed singly to prevent damage to the implanted lens. After both the stereotaxic surgeries for virus injection and GRIN lens implantation, the mice were injected intraperitoneally with 0.2ml of 1mg/ml Carprofen for 3-4 consecutive days to minimize pain.

After an additional two-week recovery period, the mice were moved to a behavior room on a 12/12-hour light/dark cycle, with lights turning off at noon and experiments conducted in the afternoon (after 2 pm).

#### Behavior

Once GRIN lens implantation was completed, mice were individually housed in a behavior room. Prior to recording, test mice underwent a habituation phase where they were connected to a dummy microscope to acclimate to its weight for 10 minutes daily over three consecutive days in their home cages. On the recording day, mice first experienced a brief 10-minute exploration period in their home cage to minimize any initial artifacts. Subsequently, recordings were conducted over several phases: a 6-minute freely moving phase, followed by a 6-minute social interaction session with an opposite-sex mouse, and another 6-minute freely moving phase. Finally, a 6-minute session involving a novel object was introduced. During the social interaction and novel object sessions, the respective stimuli (stranger mouse or object) were introduced into the home cage at 1 minute from the start of recording and remained for the subsequent 5 minutes. Between each recording session, there was at least a 10-minute interval. For repeated introductions of social or non-social cues, recordings were continuous for 8 minutes as detailed in the figures. Social or non-social cues, whether novel or familiar, were introduced at either 1 minute or 6 minutes into the recording session and removed 1 minute after their introduction. This setup allowed for the assessment of neural responses across multiple presentations of stimuli within a single recording session. To align behavior and calcium videos, a TTL pulse triggered calcium recordings through Anymaze software (Stoelting) at the start of each trial along with a behavior video recording. All Ca²⁺ signals were recorded using a miniature microscope and nVista DAQ system (Inscopix) at 20 frames per second (fps).

#### Image processing and analysis

All calcium ion (Ca²⁺) imaging movies obtained with the nVista system underwent initial preprocessing using the Inscopix Data Processing Software (IDPS; Inscopix). The GCaMP6m emission signals were captured continuously at a frame rate of 20 Hz under blue LED light (455 ± 8 nm, with power ranging from 10% to 60% and analog gain set to 1). Prior to motion correction, the spatial resolution of the videos was reduced by downsampling (4x). Following motion correction, the video data were transformed into [fluorescence (F) - background fluorescence (F₀)] / F₀ (ΔF/F₀) values, utilizing the mean projection images of the entire movie as F₀. Individual calcium signals originating from specific regions of interest (ROIs, i.e., cells) were identified using principal component analysis (PCA) and independent component analysis (ICA), as previously described(*102*). The acquired timeseries of Ca²⁺ signals (ΔF/F₀) (IDPS; Inscopix) were further analyzed and visualized using Python 3.9, NumPy, SciPy, Pandas, Matplotlib, and Seaborn.

The peaks/troughs of ΔF/F₀ were detected as follows. First, the baseline fluctuation of ΔF/F₀ (bl_fluc) or the noise level was calculated within the baseline time window (0-180sec) by averaging 5 smallest peak-trough values. This approach allowed us to exclude biological Ca^2+^ spikes in the baseline. We defined standardized ΔF/F₀, z_ ΔF/F₀, as (ΔF/F₀ - mean(ΔF/F₀)) / bl_fluc. z_ ΔF/F₀ was smoothed with Savitzky-Golay filter and peaks and troughs were detected using the functions in Python SciPy package (e.g. savgol_filter & find_peaks). | z_dF/F | >= 3 were considered as peak/trough events. The times of peaks/troughs within 0.66 sec (1/sampling rate*10) were combined and considered as a single peak/trough.

#### Data plotting and Statistical analysis

For the anterograde projections (STPT) and monosynaptic input tracing (LSFM) datasets, we used Matlab (MathWorks) and/or Prism (GraphPad) for plotting the maximum values for each anatomical area (ROI), combining data from all injections. Data was organized in Prism (GraphPad), with each individual mouse’s data included in separate columns. Maximum values for each anatomical subregion (ROI) were then computed and plotted as bar graphs. Flatmaps were generated using Matlab, while all other graphs were created using Prism version 9.

For in-vivo miniscope recordings, heatmaps showing ΔF/F₀ for individual cells were plotted using Matlab. ΔF/F₀ plots were presented as the mean ± standard error of the mean (SEM) using Prism version 9. Bar graphs showing the number of peaks were plotted using Prism after collecting data from the analysis performed using Python (as explained previously for image processing and analysis). The datasets were assessed for normality and homogeneity of variance to ensure the assumptions for parametric tests were met. For single introduction experiments (Figure 7), all datasets passed the Shapiro-Wilk normality test and were analyzed using the Wilcoxon matched-pairs signed-rank test for significance in Prism. For datasets that did not pass the normality test (Supplementary Figure 2), paired t-tests were performed to determine significance using Prism.

### Software accessibility

All custom-built codes and flatmaps used in the current study will be freely available on request and can be used without any restriction.

### Data sharing plan

Data files associated the anterograde projectome and monosynaptic input, and other experimental data will be freely available on request

## Acknowledgements

We would like to extend our sincere gratitude to all current and former members of the Yongsoo Kim Lab for their dedication, motivation, and insightful discussions, which greatly contributed to the conceptualization of this project. We thank Dr. Brian Mathur for the critical input during the manuscript preparation. Figures 1A, 2A-B, 4A, 4C, 5A-B, 7A-B, 7F, 7I, and Supplementary Figure 2A were created using BioRender.com and are licensed under a Creative Commons Attribution-NonCommercial-NoDerivs 4.0 International license. We also acknowledge the High Performance Computing cluster for providing essential computational resources, as well as the Genome Sciences Core facility for access to the MERSCOPE platform at the Penn State College of Medicine.

The Genome Sciences Core – RRID:SCR_021123 services and instruments (MERSCOPE, Agilent 2100 bioanalyzer) used in this project were funded, in part, by the Pennsylvania State University College of Medicine via the Office of the Vice Dean of Research and Graduate Students and the Pennsylvania Department of Health using Tobacco Settlement Funds (CURE). The content is solely the responsibility of the authors and does not necessarily represent the official views of the University or College of Medicine. The Pennsylvania Department of Health specifically disclaims responsibility for any analyses, interpretations or conclusions.

## Author contributions

Conceptualization: YK, SBM, SS

Data Collection and experimental support: SBM, SS, RB, AP

Data Analysis: SBM, HK, DP, DS, JKL

Code development: YW, HP, FNK

Visualization: HP

Supervision: YK

Writing: SBM, YK with help from all authors

---

**Supplementary Figure 1.**
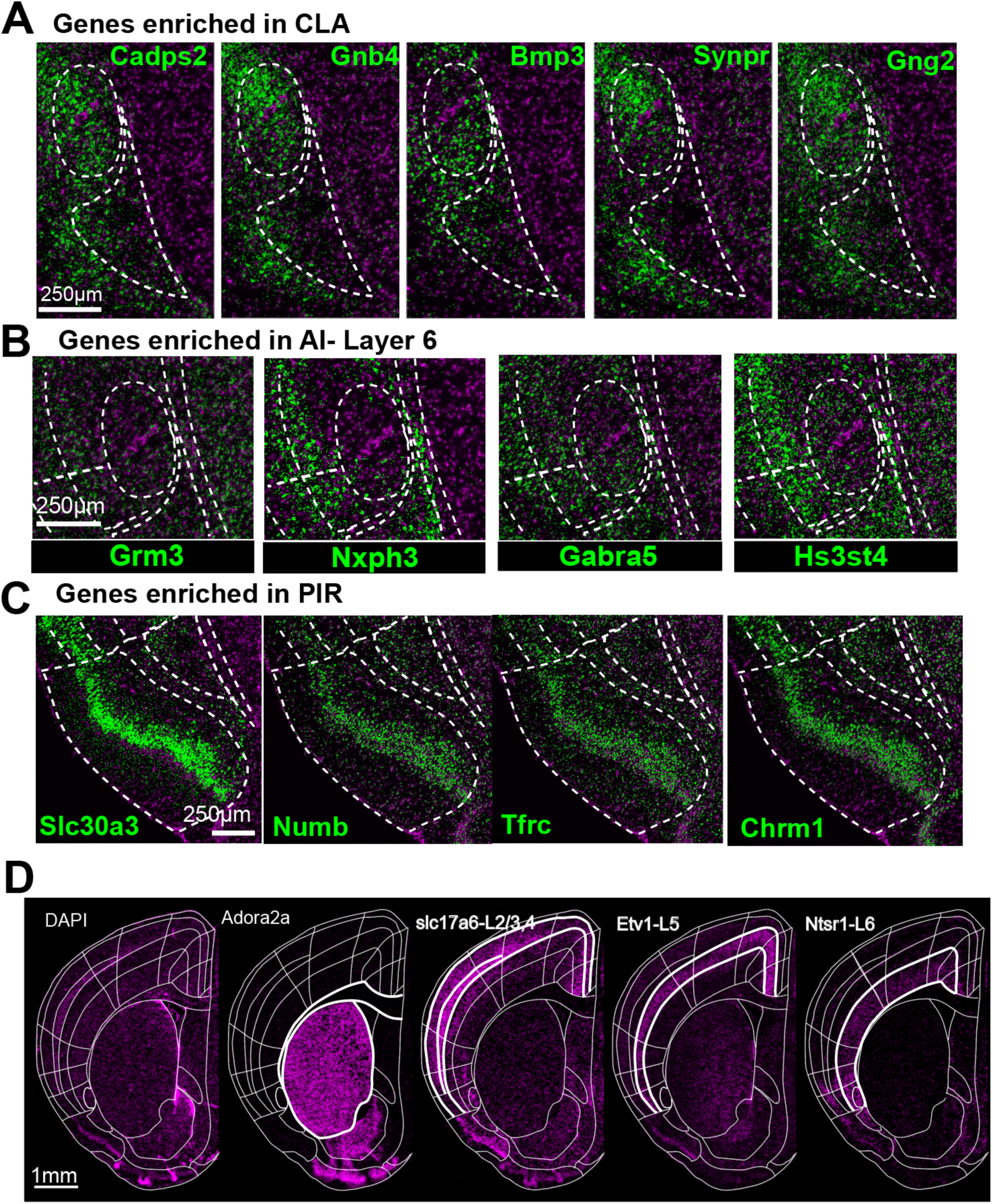
Molecular organization of the CLA, AI, and PIR. **A.** Genes enriched in the CLA (claustrum) identified through spatial transcriptomics. **B-C.** Genes enriched in AI (anterior insula) layer 6 and PIR (piriform cortex), respectively. **D.** Striatum and isocortical layer-specific markers highlighted in a spatial transcriptomics coronal section for accurate anatomical registration.

**Supplementary Figure 2.**
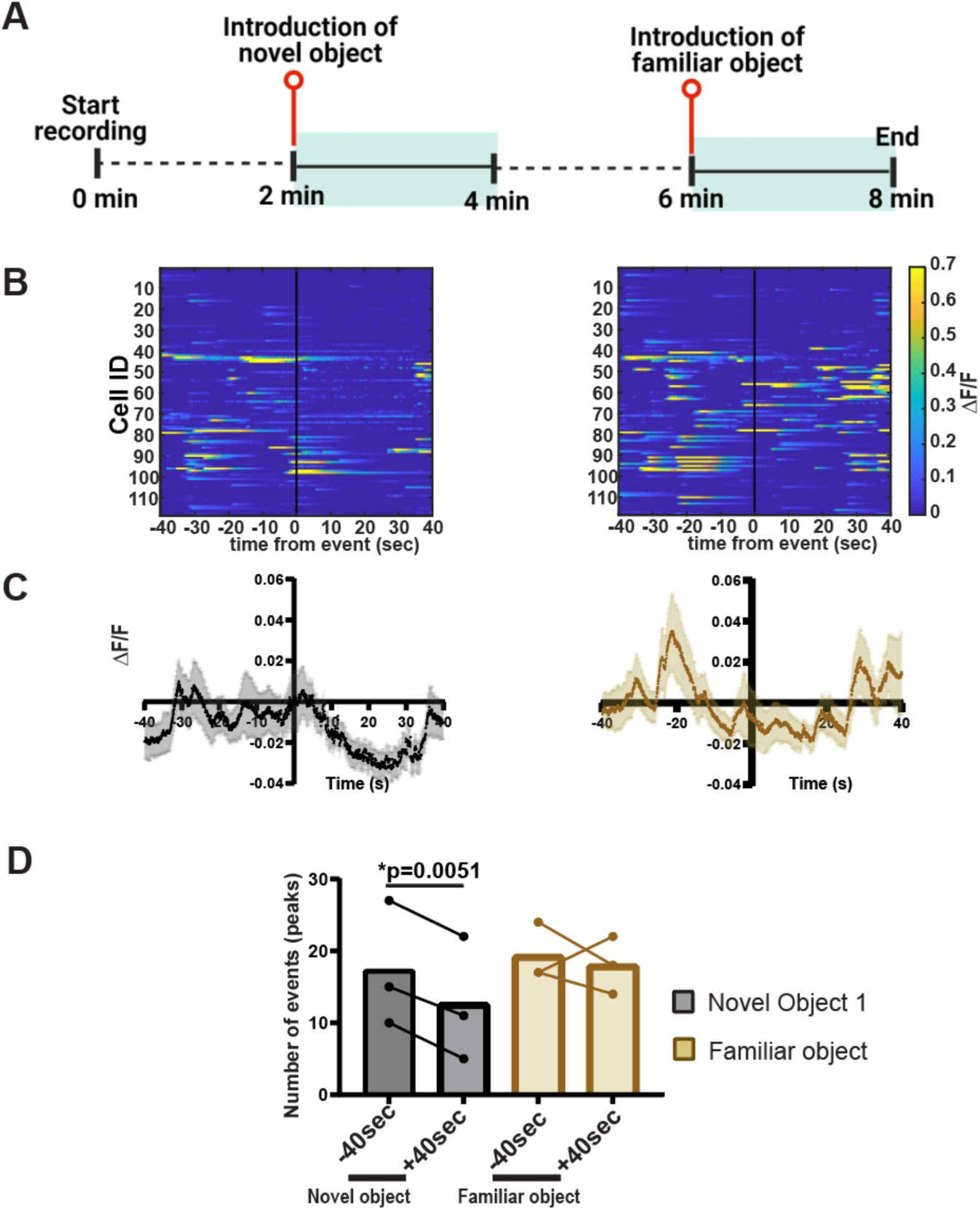
EPd neurons show reduced neuronal activity in response to non-social novel cues. **A.** Behavioral paradigm used to assess neuronal response. **B.** Heatmap displaying neuronal activity (ΔF/F) recorded 40 seconds before and after the introduction of novel or familiar objects. **C.** Comparison of neuronal activity showing a significant decrease in response to novel objects, with no significant change in response to familiar objects. **D.** Quantification of calcium events before and after the introduction of the stimulus, analyzed using the Shapiro-Wilk normality test followed by a paired t-test.

